# Cold-Shock-Mediated Inhibition of Silk Extrusion in *Galleria mellonella* (Lepidoptera: Pyralidae) for Improved Handling in Infection Studies: Effects on Developmental Traits and Pathogen Susceptibility

**DOI:** 10.64898/2026.01.20.700541

**Authors:** Deokary J. Matiya, Keira Tutt, James G. Wakefield, Jennie S. Campbell

**Author notes:** corresponding author –.

## Abstract

The greater wax moth, *Galleria mellonella*, is an increasingly important invertebrate model for infection biology, yet silk extrusion during handling complicates larval injections and hampers survival assessment. Here, we develop and characterise a simple cold-shock method that reliably inhibits silk production without compromising larval viability. Larvae exposed to −20 °C for 10 minutes completely suppress silk extrusion with 100% survival, representing a substantial improvement over previous chilling approaches. Cold-shocked larvae successfully remained capable of completing development, although pupation and adult emergence were delayed, body weight and fecundity were reduced, and wing deformities were more common. While cold-shock did not alter silk gland morphology, spinneret structure, or fibroin gene expression, confocal imaging revealed pronounced disorganisation of F-actin and α-tubulin networks within silk gland cells, indicating cytoskeletal disruption as a likely mechanism underlying silk inhibition. When challenged with *Escherichia coli*, cold-shocked larvae responded comparably to controls, with survival influenced primarily by feeding status. Together, these findings demonstrate that short-term cold-shock provides an efficient, reproducible, and easy implemented method for preventing silk extrusion in *Galleria* larvae, markedly improving handling and experimental safety while preserving their suitability as a model host for pathogen research.

**Highlights:**

- Cold-shock at −20 °C for 10 minutes inhibits silk extrusion.
- Larvae survive treatment with no loss of suitability for infection studies.
- Development slows and adult weight, fecundity, and wing quality decline.
- Silk glands stay intact; gene expression remains unchanged after cold-shock. Cytoskeletal disruption likely drives the failure of silk secretion.

## Introduction

The greater wax moth, *Galleria mellonella* (Lepidoptera: Pyralidae), is a globally distributed pest of honeybee hives and, over the past two decades, has been rapidly adopted as an invertebrate model organism in scientific research [1–3]. In particular, *Galleria* larvae are now widely used as an insect model in infection biology, antimicrobial testing and toxicity studies [4–6,2,7], and the model represents a valuable alternative to mammalian systems, supporting the 3Rs principles (Replacement, Reduction, and Refinement) [8]. Their low cost, ease of maintenance, and susceptibility to many human pathogens make them attractive to researchers and, importantly, larvae can be maintained at human physiological temperature (i.e., 37 °C), unlike *Drosophila melanogaster* and *Caenorhabditis elegans* [9]. Furthermore, their innate immune system shares broad functional similarities with mammals, and results from *Galleria* infection studies often yield results comparable to those obtained in murine models [10,11].

Despite these strengths, the majority of *Galleria* research continues to rely on larvae sourced from the pet and bait trade. These larvae are popular largely because they are easy to handle - they do not spin silk. Although silk spinning is essential to construct larval feeding tubes and to protect the developing pupae in the wild, silk production in laboratory animals can complicate pathogen injections and impede phenotypic measurements such as melanisation and mortality, while silk removal can increase the likelihood of damaging the larvae. However, the apparent convenience of pet and bait trade-supplied larvae masks significant scientific limitations. These larvae are mass-reared with no standardisation of genetic background, diet, microbial status or developmental synchrony. Their health and quality vary widely both between and within batches, and the methods suppliers use to suppress silk production are undocumented. As a consequence, researchers must often discard unfit individuals, increase sample sizes to compensate for variability, and accept a level of biological noise that would be unacceptable in any vertebrate model. This inconsistency undermines the reproducibility and interpretability of *Galleria*-based experiments and highlights the need for research-grade larvae produced under defined, controlled conditions.

Since 2021, the *Galleria mellonella* Research Centre (GMRC – www.gmrcuk.org) at the Living Systems Institute, University of Exeter, has addressed this gap by maintaining a laboratory colony of *Galleria mellonella*, supplying research-grade larvae to, and setting up sibling colonies at, institutions across the UK and Europe. However, GMRC larvae, like their wildtype counterparts, spin silk. We therefore sought to develop a standardised, reliable approach to inhibit silk spinning in the laboratory setting.

Three methods have been reported to inhibit silk extrusion in moth species: starvation, exposure to alcohol vapour, and chilling (cold-shock) [12]. Among these, cold-shock induced no visible morphological damage [12,13], whereas alcohol vapour caused detectable damage to the silk glands [14]. Starvation is generally ineffective at suppressing silk production in late-instar larvae [13], the stage typically used for infection experiments. Thus, cold-shock is considered the most promising approach, although published studies show variable effectiveness depending on species, temperature, and duration of exposure. For example, *Plodia interpunctella* larvae chilled at 4 °C for 3–12 h exhibited high silk suppression (>95%) but extremely low survival (∼5%) [15], whereas *Galleria* larvae chilled at 0 °C for 30 min to 2 hours survived well but showed only partial inhibition of silk spinning (56–77%) [16,17].

To establish an effective de-silking method based on larval cold-shock, we used a −20 °C freezer - a piece of standard equipment available in virtually all research laboratories. We systematically determined the exposure duration that maximises silk-spinning inhibition while preserving high larval survival. We also examined the morphological and subcellular effects of cold-shock on the silk gland, evaluating its impact on subsequent development, fecundity and life span. Finally, to confirm that cold-shocked larvae remain suitable for infection experiments, we compared their susceptibility with control larvae injected with a lethal dose of *E. coli*. This work therefore removes one of the major obstacles preventing wider adoption of controlled, research-grade *Galleria* lines, directly strengthening its use as a tractable, standardised model for studying host–pathogen interactions.

## Materials and Methods

### Insect Rearing

All experiments were conducted using larvae obtained from a laboratory-reared colony maintained in genetic isolation since 2017 at the *Galleria Mellonella* Research Centre (GMRC), housed within the Living Systems Institute, at the University of Exeter [6,18]. The colony was reared from eggs to adults under standard conditions for *Galleria* maintenance in *in vivo* microbiological studies. Rearing was carried out in constant darkness at 30 °C (LEEC incubator) and larvae were maintained on an artificial diet (diet recipe 3 - Jorjão et al., (2018)). The diet consisted of autolysed yeast powder (Brian Drewitt), soy flour, maize flour (both Buy Whole Foods), skimmed milk powder (Henley Bridge), glycerol (Fisher Scientific), organic honey (privately sourced), and wax pellets (Eco Lux UK).

### Larval Cold-Shock Methodology and Assessment

To determine the chilling duration that suppresses silk extrusion whilst maintaining complete survival, *Galleria* larvae were stratified into two weight categories: light (0.20–0.26 g) and heavy (0.30–0.36 g). These categories were selected to reflect the weight ranges of larvae commonly used in infection studies [20–23]. Larvae were placed at −20 °C in an underbench freezer for 5, 10, 15, and 20 minutes. At each time point, 50 larvae were placed into five Petri dishes (10 larvae per dish). The experiment was repeated once. After cold treatment, larvae were transferred to a 30 °C incubator (LEEC), and both silk extrusion and survival were monitored over 48 hours.

Based on the results, a 10-minute cold-shock duration at −20 °C was selected for further investigation. To assess the impact of cold-shock on life history traits, control and treatment (cold-shock) larvae were each placed in Petri dishes (10 larvae per dish, 6 dishes per experimental group). Both groups were provided with larval diet and maintained under standard conditions at 30 °C. Larvae were monitored throughout their development until the death of all resulting adult moths. The experiment was repeated once. Life history traits recorded included survival rate, development time, pupation and eclosion success, larval and pupal weight, fecundity, fertility, and egg hatching rate.

### Live and fixed microscopy

Control (reared at 30 °C) or heat-shocked larvae (-20 °C for 10 mins, then placed back at 30 °C for the appropriate time point) were anaesthetised using CO_2_ gas delivered via a FlyPad (FlyStuff) at room temperature (RT), before being placed in a watch glass containing Insect Physiological Saline (IPS) − 150 mM sodium chloride, 5 mM potassium chloride, 10 mM Tris HCl pH 6.9, 10 mM EDTA and 30 mM sodium citrate). Larvae were dissected under a Leica EZ4 W stereomicroscope (Leica Microsystems) at 10× magnification to isolate the silk glands. The glands were then transferred to a fresh watch galss containing IPS and examined under the same stereomicroscope. Images were captured using the stereomicroscope’s integrated 5.0-megapixel CMOS WiFi camera and transferred via WiFi Broadcast to an Android smartphone running the Leica AirLab v2.0 application, where they were viewed at 30× (zoomed out). The images were subsequently opened in FIJI — image processing software on a personal computer and examined visually to identify any gross morphological differences.

For subcellular examination, the silk glands were fixed in 4% Paraformaldehyde, diluted in PBS containing 1% Triton X-100 (PBST), for 30 minutes, immediately following dissection. Fixed glands were washed in PBST and stored at 4 °C.

To visualise nuclear morphology and the actin cytoskeleton, glands were incubated with DAPI (1:1000) and Phalloidin (1:1000) for 1 hour at RT. Subsequently, the samples were washed three times with PBST, mounted in mounting medium (glycerol: PBS, 7:3 v/v), and stored at 4 °C until imaging. Confocal imaging was performed using a Zeiss LSM 880 AiryScanner confocal at 10× (NA 0.45) and 40× (NA 1.3) magnifications.

To assess the effect of cold-shock treatment and subsequent recovery on the microtubule network, the transgenic *Galleria* line {*Gm*hsp90:GFP-αtub1b, *Bm*hsp90:histone2av-mCherry, 3xP3-dsRed2} [18] was utilised. Immediately following dissection, silk glands were placed into a 35 mm Glass Bottom Dish (No. 0, MatTek) containing IPS and imaged by confocal microscopy (Zeiss LSM 880 AiryScanner confocal, x 63, NA 1.4). All glands were imaged within 45 minutes of dissection.

### Gene Expression Analysis in the Silk Gland

To assess gene expression within the silk gland following cold-shock treatment, glands were dissected, as above, from both control larvae and treated larvae that had been returned to 30 °C for 2 hours after cold-shock. The dissected glands were manually homogenised using a plastic laboratory micropestle in 200 µL TRIzol, followed by a 5-minute incubation at RT. After adding 40 µL chloroform, samples were manually shaken for 15 seconds, incubated for 3 minutes at RT, and centrifuged at 12,000 ×g for 15 minutes at 4 °C. The upper aqueous phase was collected, and RNA precipitated with 100 µL isopropanol, incubated for 10 minutes at RT, and pelleted by centrifugation at 12,000 ×g for 10 minutes at 4°C. The pellet was washed twice with 100 μL of cold 75% ethanol, centrifuged at 12,000 × g for 5 minutes at 4 °C, air-dried for 2 minutes, and then dissolved in 50 μL of RNase-free water. RNA was incubated at 55° C for 10 minutes, quantified using a Nanodrop, and stored at –80 °C. For cDNA synthesis, 1 µL of 250 ng of RNA from each sample was combined with 1 µL of 1 pM primers, 0.5 µL of 10 mM dNTPs, and RNAse-free water to a total volume of 6 µL, heated at 65°C for 5 minutes, and then placed on ice. A premix of 2 µL 5× buffer, 1 µL DTT, 0.75 µL DEPC water, 0.05 µL RNasin (40 U/µL), and 0.2 µL M-MLV reverse transcriptase (40 U) was added. The 10 µL reaction was incubated at 37 °C for 50 minutes, then at 95 °C for 5 minutes, and subsequently purified by ethanol precipitation according to standard protocols and stored at –80 °C.

PCR was performed using the purified cDNA in 25 µL reactions using OneTaq 2× Master Mix (NEB), including 12.5 µL Master Mix, 0.5 µL each of 10 mM forward and reverse primers (0.2 µM), 1µL of cDNA from the RT reaction, and water to volume. Thermal cycling involved an initial denaturation at 94 °C for 30 seconds; followed by 35 cycles of 94 °C for 20 seconds, 56 °C for 20 seconds, and 68 °C for 1 minute per kilobase; and a final extension at 68 °C for 5 minutes. PCR products were resolved on 1% agarose gels and visualised using a gel imaging system (Bio Rad).

### Scanning electron microscope (SEM) imaging of the spinneret structure

To analyse the effect of cold-shock treatment on the structure of the larval spinneret, larval heads were dissected from control larvae, larvae immediately following 10 minutes cold-shock, and cold-shocked larvae that had been allowed to recover at 30 °C for 4 or 24 hours. Four larvae were assessed per timepoint. Prior to decapitation, larvae were anaesthetised with CO₂ as above, and the entire head was removed using a scalpel blade, cutting posterior to the pronotum. Heads were immediately fixed at room temperature in EM fixative (2% formaldehyde, 2% glutaraldehyde in 0.1 M PIPES (piperazine-N,N′-bis(2-ethanesulfonic acid) buffer) and stored at 4 °C until processing.

For SEM preparation, the fixative was removed by three washes in 0.1 M PIPES (each for 5 minutes). Heads were then post-fixed and heavy-metal stained with 1% osmium tetroxide for 1 hour, rinsed three times in distilled water, and dehydrated through a graded ethanol series (30–100%) over 1.5 hours. Samples were then incubated in 100% ethanol containing hexamethyldisilazane (HMDS) for 3 minutes, followed by a final chemical drying step in HMDS (also 3 minutes). The dried specimens were mounted on SEM stubs and coated with 10 nm gold/palladium. Imaging was performed on a Zeiss GeminiSEM 500 microscope.

### Bacterial Preparation and Larval Injection

The *E. coli* strain MG1655 was cultured in 10 mL of Luria-Bertani (LB) broth at 37 °C for 16 hr with shaking at 200 rpm. 75 μL of the overnight culture was inoculated into 10 mL of fresh LB and incubated for an additional 2 hours under the same conditions. The culture was then transferred to 1.5 mL centrifuge tubes and centrifuged at 5,000 ×g for 5 minutes. After discarding the supernatant, the bacterial pellets were weighed and suspended in phosphate-buffered saline (PBS) to a final concentration of 70 mg/mL. The suspension was vortexed thoroughly, and the bacterial density was measured at an optical density of 600 nm (OD_600_), yielding an average reading of 1.6 for a 10× bacterial dilution, which corresponded to approximately 1.28 × 10¹⁰ cells/mL.

*Galleria* larvae (0.30–0.36 g), both cold-shock and control individuals. Cold-shock larvae were pre-treated by placing them at –20 °C for 10 minutes, then returned to a 30 °C incubator until injection, which was performed either 2 or 4 days post cold shock, depending on the experimental group. Larvae were further subdivided into three treatment conditions: fed for 4 days post-chilling, or starved for either 2 or 4 days. Each subgroup contained 60 larvae, housed in petri dishes in groups of 10. Control groups received PBS injections, while experimental groups received injections of the prepared *E. coli* suspension. A total of 1,800 larvae were used: 700 cold-shock-infected, 700 control-infected, 200 cold-shock-PBS, and 200 control-PBS.

Each experimental larva was injected with 10 µL of the bacterial suspension (1.28 × 10⁸ cells/mL) into the last right proleg using a Hamilton 700 series syringe (Merck) fitted with a PB600 repeating dispenser. The control groups, comprising both cold-shocked and untreated larvae, were each injected with 10 µL of PBS. Post-injection, larvae were incubated at 37 °C, and survival was assessed at 24 and 48 hr.

### Data analysis

Statistical analyses were performed using IBM SPSS Statistics version 22. Normality was assessed with the Shapiro–Wilk test, and group comparisons for continuous variables (e.g., weights, fecundity, hatching rate) were conducted using unpaired two-tailed Student’s t-tests. Categorical outcomes (e.g., pupation, survival) were analysed using χ² tests and Kaplan–Meier survival analysis. Survival following *E. coli* infection was assessed using Kaplan–Meier survival analysis, with group differences evaluated using the log-rank test. A significance threshold of p < 0.05 was used throughout the data analysis. Box plots with annotations indicating statistical differences between groups and Kaplan–Meier survival curves were generated in SPSS to visualise the data.

## Results

### Establishing consistent parameters for cold-shock-mediated inhibition of silk extrusion

We began by investigating the shortest duration of cold-shock at −20 °C that could effectively suppress silk extrusion in *Galleria* larvae while maintaining survival. In our initial experiments, we stratified larvae into two weight categories: light (0.20–0.26 g) and heavy (0.30–0.36 g), reflecting the weight ranges of larvae commonly used in susceptibility studies [20–23]. We found that exposure for 5 or 10 minutes resulted in 100% survival across both weight ranges (0.20–0.26 g and 0.30–0.36 g; Fig. 1A). However, complete inhibition of silk extrusion was achieved following a minimum cold-shock duration of 10 minutes (Fig. 1B). At 15 minutes, survival declined to 73% in lighter larvae (range, 60–80%) and 86% in heavier larvae (range, 70–90%), with a significant difference between groups (t_(17)_ = –3.74, p = 0.002). At 20 minutes, survival dropped further to 23% in lighter larvae (range, 10–40%) and 40% in heavier larvae (range, 30–60%) (t_(19)_ = –3.2, p = 0.004). Thus, 10 minutes of cold-shock at −20 °C was determined to be optimal, fully inhibiting silk extrusion while preserving maximum survival, irrespective of larval weight.

**Figure 1.**
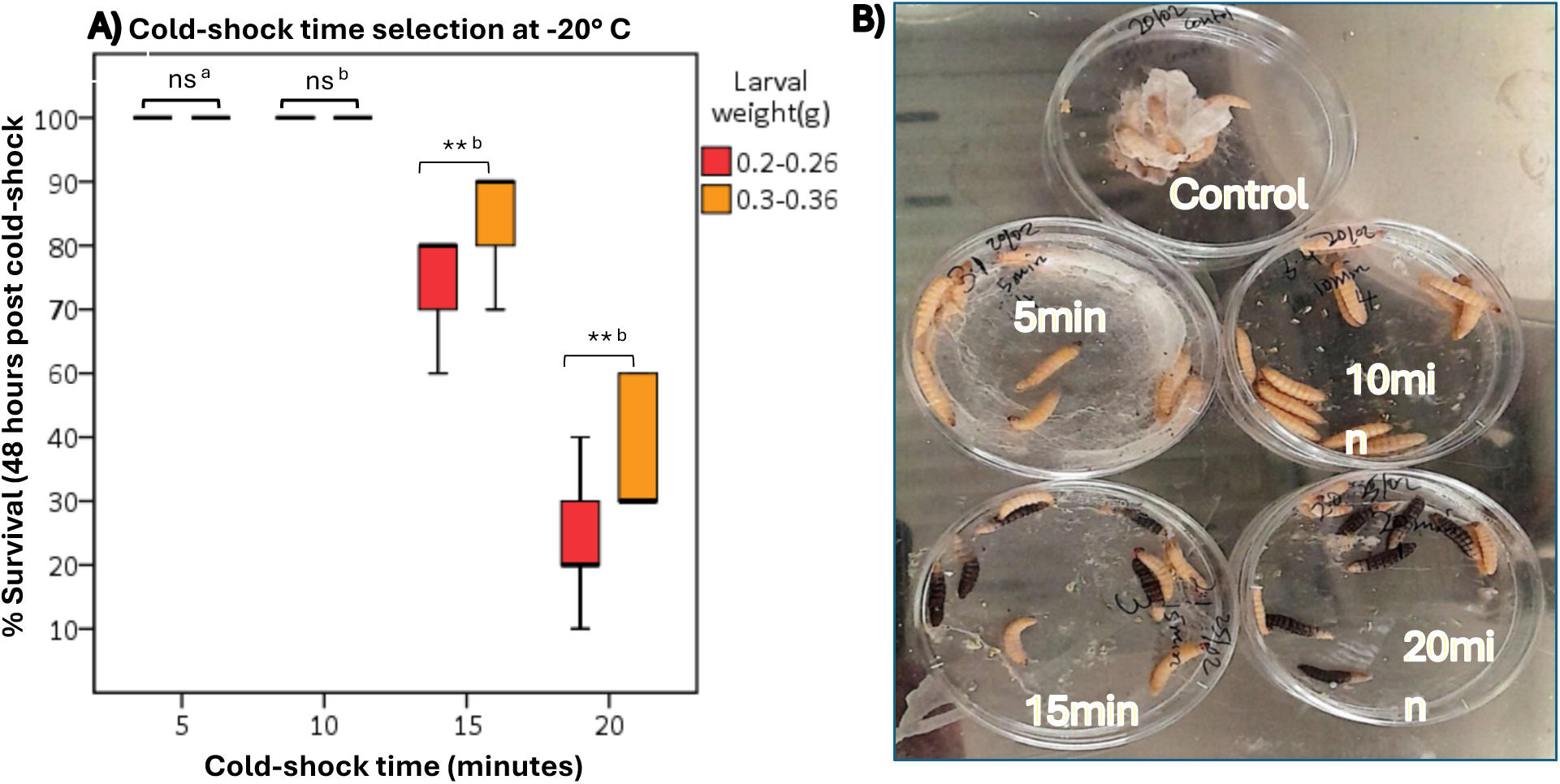
Effect of −20 °C cold-shock duration on silk extrusion and larval survival in *Galleria mellonella*. (A) Larval survival (%) at 24 and 48 hr post–cold-shock across different exposure times and weights. (B) Representative images of larvae after treatments; black larvae indicate dead individuals and whitish material indicates silk tubes. Key: * = p < 0.05; ns = p > 0.05; a = extruding silk; b = not extruding silk.

### The effect of cold-shock on *Galleria* larval and pupal weight, development time and survival rates

To characterise the effects of cold-shock treatment on larval and pupal development, beyond the loss of silk extrusion in *Galleria mellonella*, treated larvae were returned to 30 °C in the presence of a food source. Cold-shock significantly influenced development time, survival, and weight (Fig. 2). Although both larval and pupal weights decreased over time in the cold-shock and control groups, the reduction was more pronounced in cold-shocked individuals (Fig. 2A). In the cold-shock group, mean larval weight averaged 0.270 g across the larval period, decreasing from 0.350 g at the start to 0.220 g by the end. The weight of cold-shocked larvae was significantly lower than that of the control group, which averaged 0.312 g and declined from 0.350 g to 0.270 g (t₍₈₃₎ = –6.9, p < 0.001). A similar pattern was observed during the pupal period. Cold-shocked individuals exhibited a mean pupal weight of 0.194 g, decreasing from 0.260 g to 0.130 g, whereas controls averaged 0.231 g (range: 0.210–0.260 g), declining from 0.260 g to 0.231 g (t₍₁₁₎ = 129.9, p < 0.0001).

**Figure 2.**
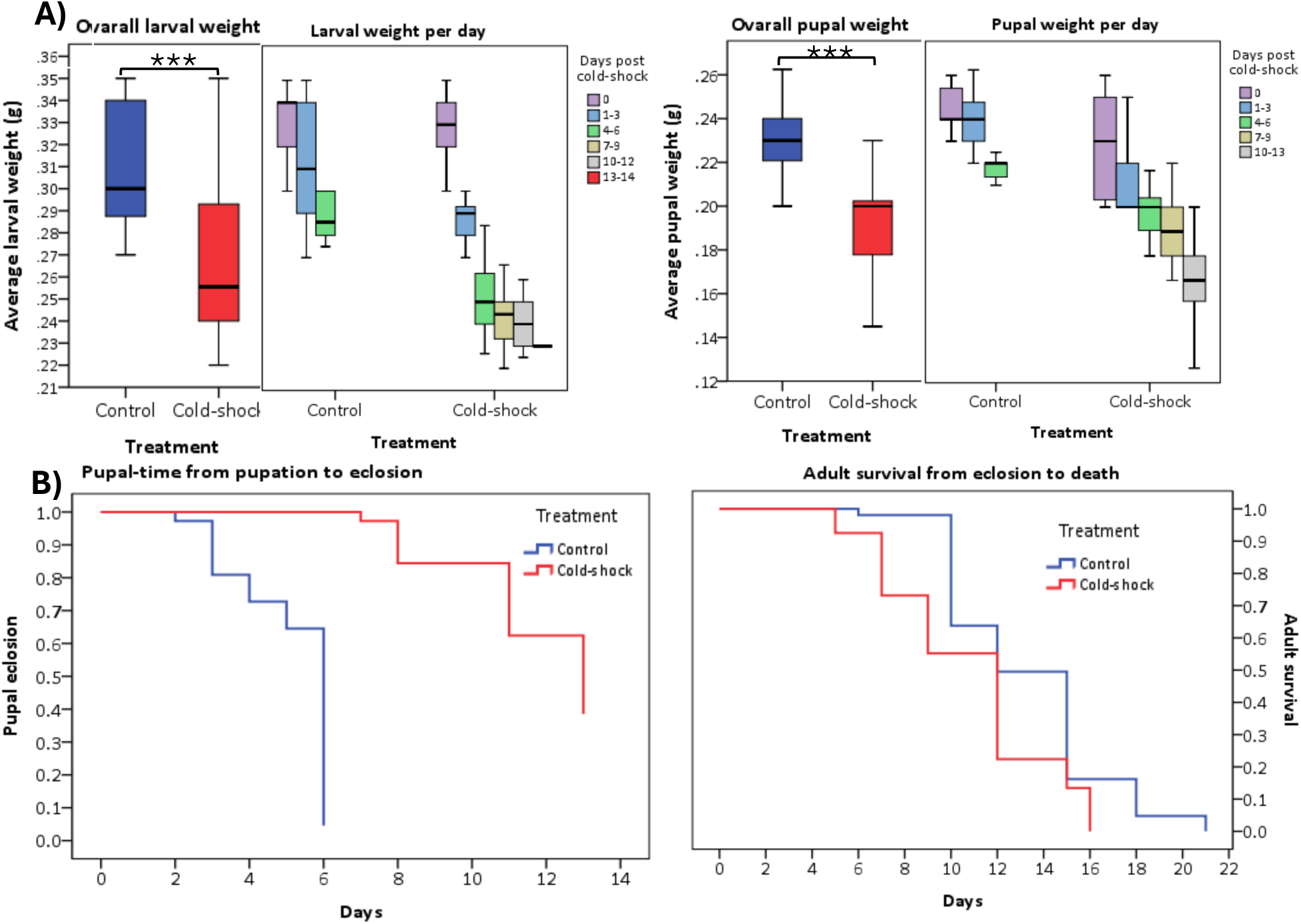
Cold-shock effects on *Galleria mellonella* larvae and pupae. (A) Box plots of overall and daily larval and pupal weights in control and cold-shock treatments. (B) Survival curves showing larval duration from cold-shock to pupation and pupal time from pupation to eclosion. Key: *** = p < 0.001 (significant); ns = p > 0.05 (not significant).

With respect to development and survival, cold-shock extended both larval and pupal durations while reducing survival rates (Fig. 2B). Control larvae pupated in a mean of 5.3 days, with 100% pupation achieved by day 7. In contrast, cold-shocked larvae required 9 days, with only 90% pupating by day 14 (χ^²^ _(1)_ = 171.7, p < 0.0001). Similarly, control pupae eclosed in a mean of 5.2 days with 96% success by day 8, compared to 11.8 days and 61.5% eclosion by day 13 in pupae developed from cold-shocked larvae (χ² _(1)_ = 192.2, p < 0.0001). These findings indicate that cold-shock delays development, lowers survival, and reduces adult longevity.

### Cold-shock effects on adult morphology, fecundity, and offspring viability

Upon eclosion of imagoes from their pupal cases, we observed physical deformities – most notably of the wing – in both adult males and females that have been exposed to cold-shock as larvae (Fig. 3A). The incidence of wing deformities was significantly higher in adults grown from cold-shock larvae (25%; n = 30/120) compared to those from control larvae (1.7%; n = 2/120).

**Figure 3.**
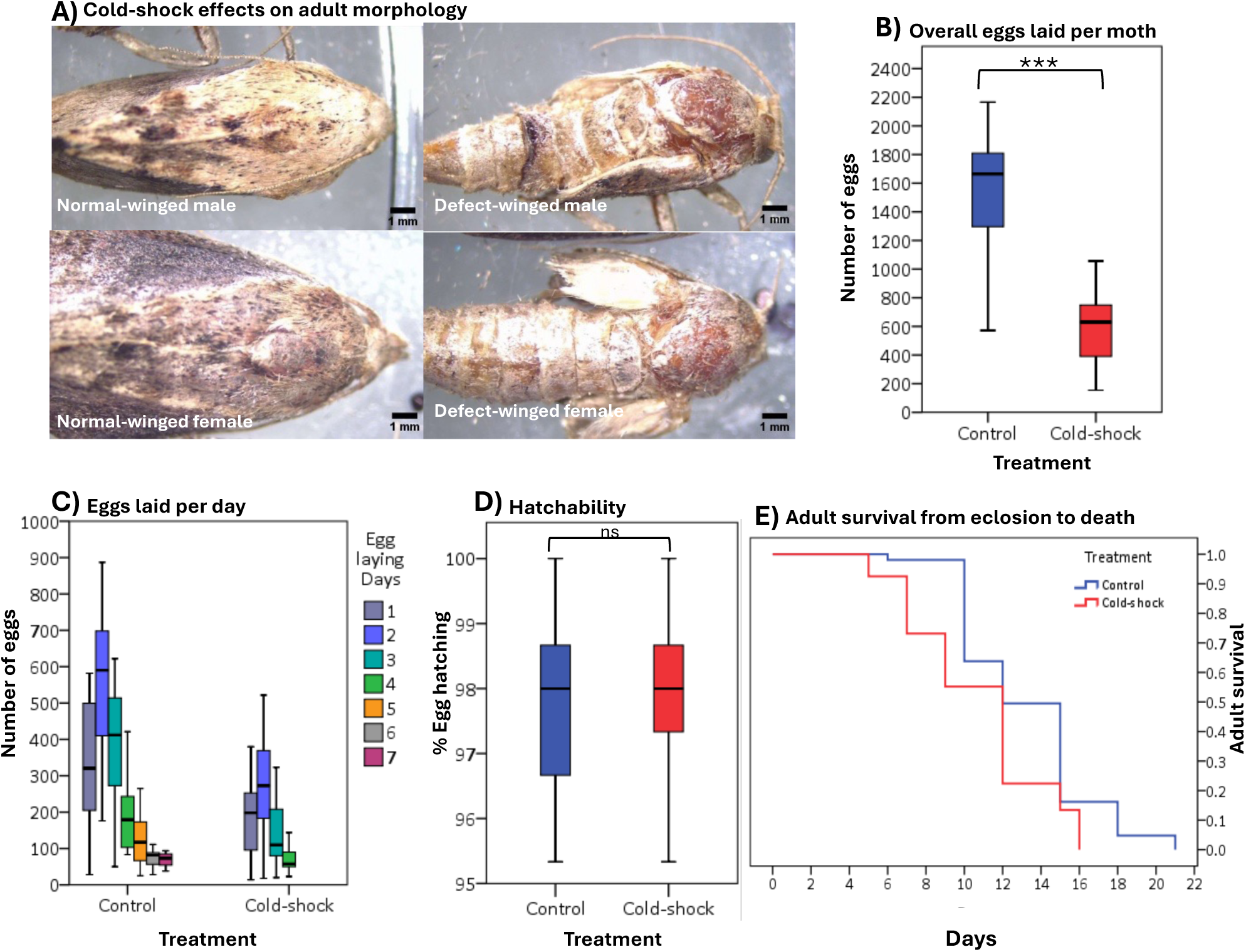
Cold-shock effects on adult *Galleria mellonella*. (A) Images showing normal- and defect-winged adults. (B) Average number of eggs laid over the moth’s lifespan.(C) average number of eggs laid per day, (D) average egg hatching rate, and (E) Adult lifespan illustrated by survival curves for control and cold-shock treatments average egg hatching rate. Key: *** = p ˂ 0.001(significant) and ns = p > 0.05 (not significant).

To assess whether larval cold-shock affected imago reproductive parameters, oviposition and subsequent hatchability were observed (Fig. 3). Cold-shock-treated females laid fewer eggs, averaging 578 per individual (range: 154–1054), while control females laid 1542 (range: 572–2166) (*t*_(31)_ = –8.59, *p* < 0.0001)(Fig. 3B). The egg-laying period lasted 4 days in the treated group and 7 days in the control group (Fig. 3C). In both cases, peak oviposition occurred on day 2, followed by a sharper decline in the cold-shock-treated group. Despite these effects, egg viability remained unaffected. Hatching rates were 97.8% (range: 97.0–98.6%) in the treated group and 98.2% (range: 97.5–99.0%) in the control group, with no statistical difference observed (*t*_(40)_ = –0.052, *p* = 0.95) (Fig. 3D).

Finally, we assessed life-span. Adult lifespan averaged 13.3 days for control moths, with 22% surviving to day 21, versus 10.7 days and 0% survival past day 17 for treated adults (χ²_(1)_ = 19.4, p < 0.0001) (Fig. 3E).

### Cold-shock treatment does not alter silk gland morphology or fibroin gene expression

Having characterised the broader physiological and developmental effects of cold-shock on *Galleria* larvae, we sought to determine the underlying mechanism of silk inhibition following treatment. To investigate the effect of cold-shock treatment on overall silk gland morphology, glands were dissected out from larvae immediately following treatment and visually compared to control glands. We found no overt morphological difference (Fig. 4A) – glands maintained their characteristic tubular structure and size, and remained glassy and translucent in appearance, suggesting that silk production is not inhibited due to gross morphological defects during silk gland development.

**Figure 4.**
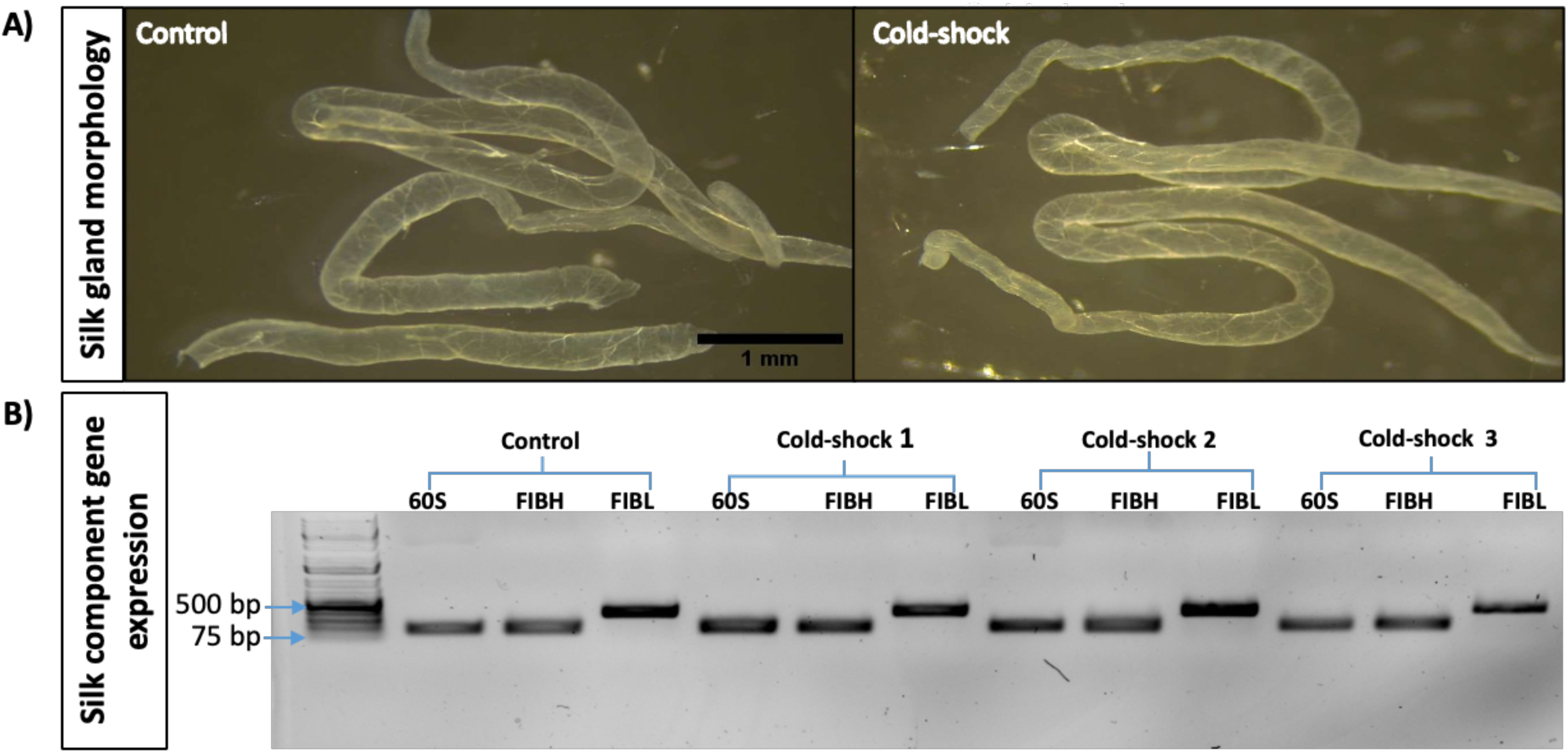
Effects of cold-shock on *Galleria mellonella* larvae. (A) Silk gland morphology in control and cold-shock–treated larvae. (B) Gel electrophoresis showing expression of silk-encoding genes (FIBH, FIBL) in control and cold-shock–treated larvae; 60S was used as the positive control, and cold-shock 1–3 indicate biological replicates.

We next examined whether silk production was disrupted at the molecular level. *Galleria mellonella* silk is mainly composed of Fibroin heavy (FIBH) and light chains (FIBL) [24]; therefore, we examined whether cold-shock treatment at −20 °C affected the expression of these major components. Using RT-PCR on mRNA extracted from silk glands two hours post-treatment, we compared transcript levels in cold-shocked and control larvae. Both Fibroin heavy and light chain transcripts remained present post cold-shock (Fig. 4B), suggesting that alteration in the levels of gene expression of the primary silk components is not responsible for the cessation of silking, post-cold-shock.

### The larval spinneret remains intact following cold-shock treatment

As silk gland morphology and fibroin gene expression appeared unaffected by cold shock, we next investigated whether structural disruption of the silk extrusion apparatus (the spinneret) might explain the observed inhibition of silk production. To assess this, we performed SEM examinations of spinnerets dissected from larval heads immediately after cold-shock treatment (t = 0) as well as at 4 and 24 hours post-treatment. At all-time points, we observed no structural abnormalities in the spinneret (Fig. 5), nor the surrounding mouthparts and cuticle. The structure remained intact, with a clearly visible opening in all samples (n = 4 per condition). In control larvae, a silk strand was visible extending from the spinneret, consistent with active silk production. Thus we conclude that spinneret damage or external blockage are not the mechanisms by which silk production is inhibited following cold-shock treatment.

**Figure 5.**
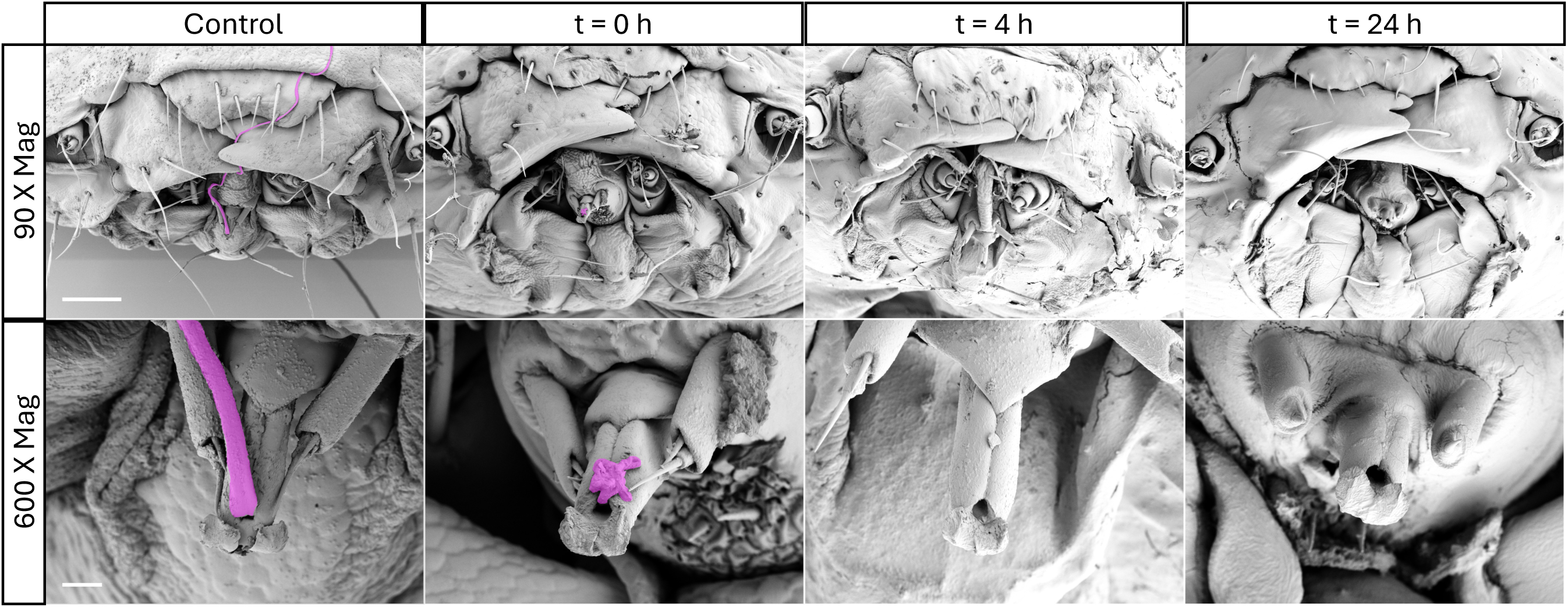
SEM micrographs showing intact spinneret structure after cold-shock treatment. Silk thread pseudocoloured in magenta (50% opacity), visible in controls, residual thread at t = 0 h, and absent at 4 h and 24 h. Scale bars: 90 X panels = 200 µm, 600 X panels = 20 µm.

### Alterations to silk-gland cellular architecture following cold-shock treatment

With no detectable abnormalities in silk gland morphology, fibroin gene expression, or spinneret structure, we examined whether cold-shock affected silk production at the cellular level. As disruption of the cytoskeleton could feasibly affect silk fibre formation, given its essential role in the intracellular transport and release of secretory cargo into the silk gland lumen, we assessed the integrity of nuclear and cytoskeletal architecture in silk gland epithelial cells. Nuclear architecture appeared normal in fixed and stained preparations of the posterior silk gland (PSG), with polyploid nuclei maintaining their typical morphology and showing no evidence of condensation or fragmentation. However, cytoskeletal organisation exhibited clear temperature-dependent alterations.

In fixed samples of the middle silk gland (MSG), Phalloidin staining revealed bright, thread-like F-actin aggregations indicative of increased actin bundling, which persisted throughout the 24-hour recovery period following cold shock (Fig. 6A). By contrast, the microtubule network was assessed using the transgenic *Galleria* line expressing live GFP–αTub1b, allowing visualisation of dynamic cytoskeletal responses immediately after dissection. In the PSG, α-tubulin fluorescence intensity was markedly reduced at t = 0 h, consistent with rapid microtubule depolymerisation. Although the network partially recovered between 4 h and 24 h post-treatment, re-establishing filamentous structures, the microtubules remained more diffuse and disorganised than in untreated controls, suggesting that cold shock induced a lasting disruption to microtubule integrity (Fig. 6B).

**Figure 6.**
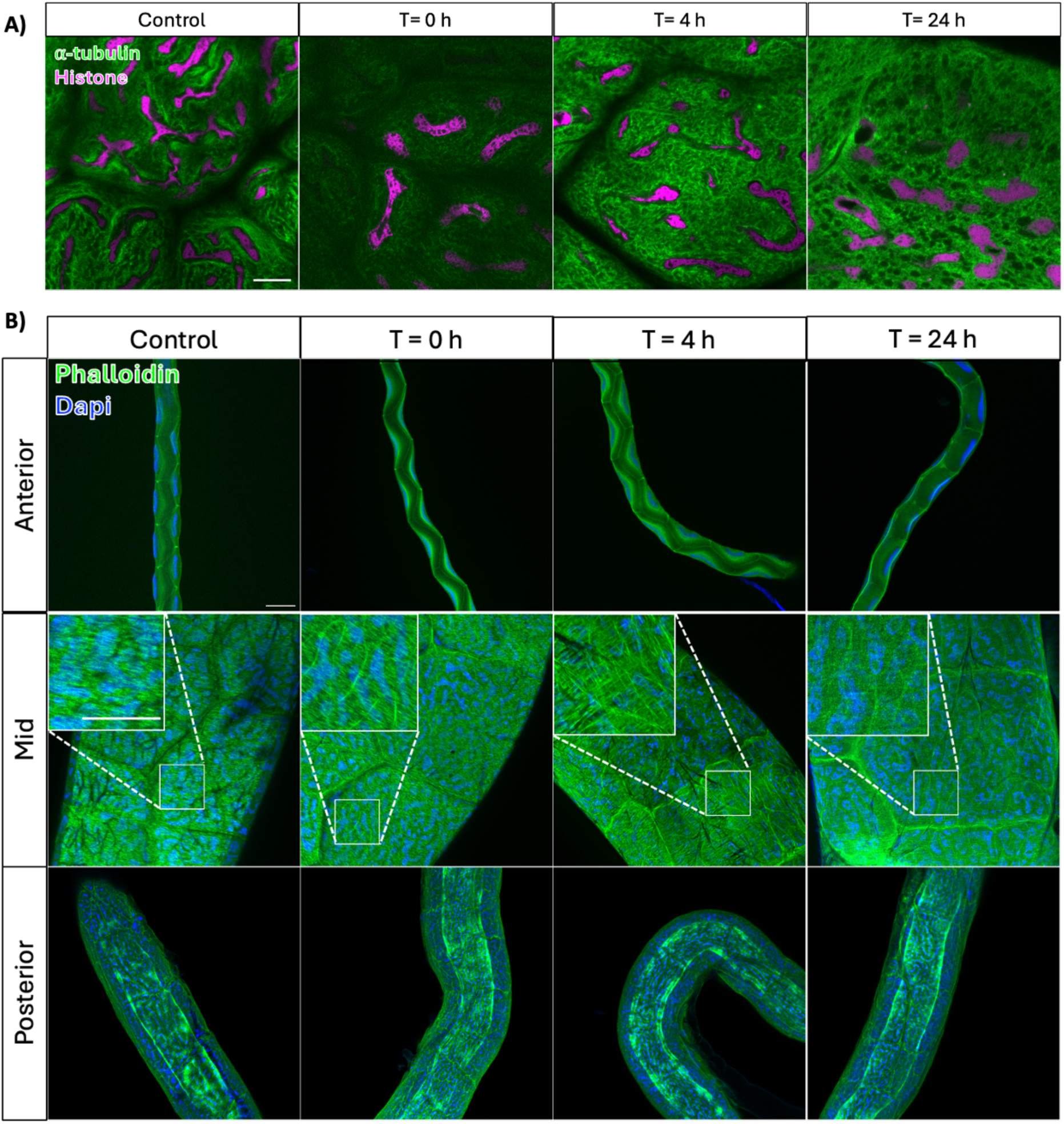
Confocal images showing the effect of cold shock on silk gland cells of *Galleria mellonella*. (A) Sections of the silk gland stained for F-actin with phalloidin (green) and nuclei with DAPI (blue). All scale bars = 100 µm. (B) Posterior silk gland of transgenic larvae expressing α-tubulin (green) and histone–mCherry fusion proteins (magenta) to visualise nuclei. Scale bar = 20 µm.

### Survival Response to *E. coli* Infection Following Chilling Under Varying Nutritional and Weight Conditions

To confirm the suitability of cold-shock larvae for infection studies, we compared the susceptibility of cold-shock–treated and control larvae to a lethal dose of *E. coli*, under both fed and starved conditions (Figure 7). Batches of control or cold-shocked larvae were returned to diet or to petri dishes without diet, for 4 days prior to inoculation with ∼1.26 x 10^6^ *E.coli* cells. Those in the cold-shock group showed significantly lower survival compared to control larvae (38% vs 59% at 24 h and 12% vs 29% at 48 h; χ²_(3)_ = 13.1, p < 0.0001). In contrast, starved larvae showed higher survival following cold-shock than in the control group (40% vs 17% at 24 h and 8% vs 3% at 48 h; χ²₍₃₎ = 22.8, p < 0.0001). When larvae were starved for 2 days, survival did not differ between cold-shock and control groups (48% vs 46% at 24 h and 16% vs 13% at 48 h; χ²₍₃₎ = 0.24, p = 0.62).

**Figure 7.**
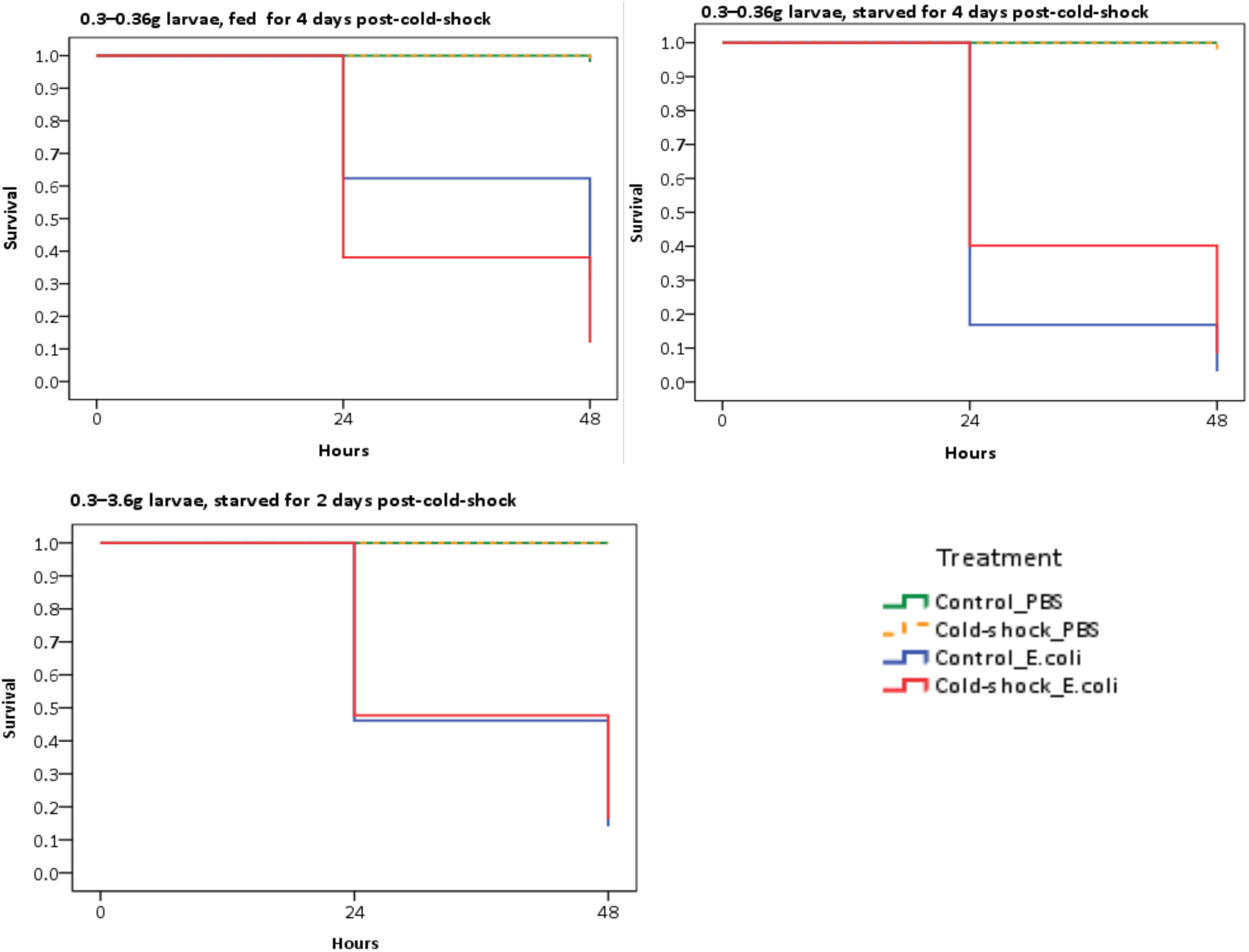
Survival of cold-shock and control *Galleria mellonella* larvae with varying body weights and feeding statuses following infection with a lethal dose of *E. coli* strain MG1655. The results are presented in survival curves.

## Discussion

In our efforts to improve the handling of *Galleria mellonella* larvae by inhibiting silk extrusion, we focused on overcoming the practical challenges that commonly arise during infection studies. Silk can increase contamination risk and necessitate manual de-silking when working with virulent microorganisms. We found that a brief cold-shock at −20 °C for 10 minutes completely prevented silk extrusion while maintaining 100% larval survival. This represents a very notable improvement over previous work in which chilling at 0 °C for 30 minutes to 2 hours produced only partial inhibition of larval silk spinning (56–77%) despite high survival [16,17]. Our approach is also highly practical: it requires only a standard −20°C freezer, routinely available in most laboratories. Thus, this simple and accessible method provides an efficient, reproducible, and safer means of preparing larvae for infection assays without compromising viability.

To assess the broader implications of cold-shock, we examined its effects on life history traits to determine the suitability of treated individuals for colony establishment and maintenance. Cold-shocked larvae completed development, and resulting females produced viable eggs, indicating that the life cycle remains intact. However, compared with controls, cold-shock prolonged both larval and pupal development, reduced body mass, decreased adult emergence, shortened lifespan, increased wing deformities, and lowered fecundity. Similar delays in development have been reported previously in chilled *Galleria* larvae [25,17,26,27] and extended pupal duration has also been documented in other moth species following low temperature exposure [15].

These effects may result from cold-induced elevation of juvenile hormone or damage to the salivary glands, both of which can disrupt normal growth and metamorphosis [17,26]. Pupal desiccation caused by inhibited cocoon silk spinning [28], together with prolonged pupal duration, as also observed in this study, may explain the low emergence and high frequency of wing deformities observed under cold shock. Overall, although cold-shocked larvae are well suited for infection studies, their altered developmental and reproductive traits make them less appropriate for long-term colony maintenance and may, at least in part, explain anecdotal difficulties in establishing colonies from pet shop larvae.

To determine whether silk gland damage underpinned the inhibition of silk extrusion, we examined the morphology and subcellular organisation of the silk gland following cold shock. Cold exposure did not produce detectable morphological alterations in either the silk gland or the spinneret compared with controls - in contrast with previous reports that described solidification of the silk gland following cold exposure [15,17]. This difference is likely due to the shorter cold exposure used in our experiment, which may have minimised major tissue damage. Indeed, we found nuclear integrity to be preserved and expression of the major silk protein gene, fibroin, unaffected after our 10 minute treatment. The resilience of these cells may reflect the polyploid nature of silk gland cells, which are known to be resistant to low-temperature stress [29].

Having established that the silk gland architecture and fibroin expression were unaffected, we next assessed the intra-cellular cytoskeletal organisation central to protein secretion. Specifically, microtubules, which direct vesicular transport, and filamentous actin (F-actin), which maintains cell morphology and supports exocytosis [30,31]. Cold-shock induced conspicuous reinforcement of F-actin networks concurrently with the collapse of the microtubule network. Strengthened F-actin may represent a compensatory response to maintain cellular integrity or enhance membrane stability under stress, whereas the disassembly of microtubules likely disrupts intracellular transport and secretion of fibroin and sericin. Comparable cytoskeletal disruptions under cold stress have been reported in several insect species [32–34]. In our study, the alterations persisted after larvae were returned to normal incubation temperatures, suggesting a lasting structural impact. This aligns with previous findings in the malt fly *Chymomyza costata*, where cold-induced microtubule disorganisation in fat body cells remained permanent without impairing survival [33]. Collectively, these results indicate that cold shock disrupts cytoskeletal integrity in the silk gland, hindering intracellular transport and secretion, thereby inhibiting silk extrusion - consistent with observations in *Bombyx mori*, where chemical cytoskeletal disruption similarly stops fibroin transport and release into the lumen [30,31].

We also assessed susceptibility to bacterial infection by comparing cold-shocked and control *Galleria* larvae challenged with a lethal dose of *E. coli*. Cold-shocked larvae remained suitable for infection studies, though survival outcomes were influenced by feeding status and timing of infection post-treatment. Fed larvae challenged four days after cold-shock showed reduced survival compared with control, possibly because cold-induced stress slowed feeding and temporarily limited energy reserves for immune responses. In contrast, cold-shocked, starved larvae exhibited higher survival than controls, possibly because control larvae expend energy on silk production while cold-shocked larvae do not. This suggests that the metabolic savings associated with inhibited silk spinning may enhance resilience under nutritional stress - consistent with the faster weight loss observed in starved control larvae compared with their cold-shocked counterparts.

Importantly, our method resolves a long-standing trade-off in the field: researchers must typically choose between non-silking larvae from pet or bait shops - whose health status is substantially compromised and life history and genetics unknown - or lab-reared, research-grade larvae, that continue to produce silk. Silk production can be off-putting to microbiologists, impeding handling, delaying injections, increasing the risk of accidental needle sticks when using virulent pathogens and complicating mortality scoring, particularly when larvae are concealed within silk or suffer damage during removal. Our simple method for generating healthy, non-silking, research-grade larvae, therefore, has considerable practical value and aligns with 3Rs principles. Moreover, because cold-shocked larvae remain suitable for infection studies for several days without feeding, they can be transported with ease between research sites. This raises the possibility of maintaining a small number of research-grade *Galleria* colonies at regional hubs, supplying high-quality larvae to multiple laboratories and thereby transforming their use in *in vivo* infection biology and AMR research.

## Notes

### Competing Interest Statement

The authors declare no competing interests. J. S. Campbell and J. G. Wakefield are co-directors of the Galleria mellonella Research Centre (GMRC), an internal research centre at the University of Exeter. The work was conducted as an academic research project.

